# Profilin-1 Promotes Chromophobe Renal Cell Carcinoma Malignancy

**DOI:** 10.64898/2026.05.11.723846

**Authors:** Kaylee Montanari, Anup Acharya, Cais Vo, Dhwani Shah, Elizabeth Henske, David Gau

**Affiliations:** Department of Pathology, University of Pittsburgh; Department of Bioengineering, University of Pittsburgh; Department of Biology, University of Pittsburgh; Brigham and Women’s Hospital, Harvard Medical School

## Abstract

Chromophobe renal cell carcinoma (ChRCC) accounts for 5% of all renal cancer cases. Despite its generally indolent behavior and low mutational burden, there is no targeted therapy for metastatic ChRCC. Profilin-1 (Pfn1), a cytoskeletal regulator of actin and tubulin dynamics, has emerged as a potential oncogenic driver in several cancers including RCC, but its role in ChRCC, remains undefined. We observed elevated Pfn1 expression in stage IV ChRCC patients, implicating Pfn1 in advanced disease progression. To investigate this, we manipulated Pfn1 expressions in two ChRCC cell lines UOK276 and RCJ41M. Pfn1 knockdown (KD) significantly reduced proliferation, invasion, and colony formation, whereas Pfn1 overexpression (OE) in UOK276 enhanced ChRCC aggressive phenotypes. Pharmacological inhibition of Pfn1 significantly suppressed proliferation and clonogenic growth in both cell lines. Additionally, Pfn1 KD increased intracellular ROS accumulation, while overexpressed reduced ROS levels, linking cytoskeletal regulation to oxidative stress control. Together, these findings position Pfn1 as a critical mediator of ChRCC progression, linking cytoskeletal remodeling to aggressive tumor behavior. This work highlights Pfn1 as a potential therapeutic target and establishes a framework for cytoskeletal-focused strategies in advanced ChRCC.

## INTRODUCTION

The American Cancer Society estimated in the United States there would be 80,000 new cases of kidney cancer in 2025 and around 14,000 deaths from the disease^1^. Chromophobe renal cell carcinoma (ChRCC) is the second most common non-clear cell renal cell carcinoma. ChRCC is thought to arise from the mitochondria-rich intercalated cells distal nephron^2^ ChRCC harbors abundant, albeit dysfunctional mitochondria, designating a unique pathological characteristic from which ChRCC differs from other RCCs^3,4^. ChRCC is more frequent in women (unlike clear cell renal cell carcinoma), associated with whole chromosome loss, typically chromosomes 1, 2, 6, 10, 13, 17, 21, and has a low mutational burden^5^. Birt-Hogg Dube (BHD) and tuberous sclerosis complex (TSC), two hereditary cancer syndromes, are typically associated with ChRCC^3^. While ChRCC is generally considered a less metastatic cancer and often has a favorable prognosis, patients with advanced stage ChRCC have a 2-year median survival rate with strikingly inadequate therapeutic options^6^. Treatment routes are limited and often repurposed from other renal cancers; immune checkpoint inhibitors, mammalian target of rapamycin (mTOR) inhibitors, and vascular endothelial growth factor (VEGF) inhibitors have demonstrated minimal response rates in ChRCC patients^7^.

The actin cytoskeleton is an important biochemical and biomechanical regulator of cancer progression across essentially all cancer types^8^. The cytoskeleton plays diverse roles in cancer can be broadly categorized into functions within cancer cells, interactions with other cells, and regulation of cancer-extracellular matrix (ECM) ^9,10^. These roles make the actin cytoskeleton a promising target for understanding and manipulating cancer behavior. One such important regulator of actin cytoskeleton is Profilin1 (Pfn1). Pfn1 is an actin binding protein and major regulator of actin dynamics, facilitates actin polymerization by promoting the exchange of ADP to ATP, regulating filament assembly. Pfn1 enables cells to form lamellipodia, filipodia, and other cytoskeletal structures necessary for migration and invasion^11,12^. Pfn1 has cancer-type-specific roles, functioning as a tumor-suppressor in some cancers like breast cancer,^13^ while demonstrating tumor permissive roles in non-small cell lung cancer, lung adenocarcinoma, gastric cancer, and ccRCC^14,15,16–19^. Studies prove and continue to explore how the diverse effects of Pfn1-actin regulation are finely tuned to the precise cellular and microenvironmental context of specific tumors.

We previously reported that, interestingly, Pfn1 mRNA expression was found generally lower in ChRCC compared to clear cell and papillary renal cell carcinoma via TCGA-GEPIA analysis^20^. Prompted by Pfn1’s promotion of tumorigenesis in RCC^19^, it’s unexplored axis in ChRCC, and the seemingly opposite expression in ChRCC, we began to explore how Pfn1 impacts ChRCC progression. In this study, we use *in vitro* functional tumor assays and multiple ChRCC cell lines to elucidate the impact of Pfn1 on ChRCC.

## METHODS

### Cell culture

UOK276 cells, RCJ41M, and RCJ41T2 cells were cultured in DMEM, 10% fetal bovine serum (FBS) and antibiotics (100 units/ml penicillin and 100 µg/ml streptomycin). UOK276 cells were supplemented with 0.05% GlutaMAX (ThermoFisher Cat: 35050079) and RCJ41M and RCJ41T2 cells were supplemented with 1% HEPES, and 1% GlutaMAX. Cells were maintained in 37°C and at 5% CO_2_.

### Pfn1 knockdown and overexpression

Cells were plated at 2×10^5^ cells/well in a 6 well plate and were transfected with either smart-pool scrambled control or Pfn1 siRNA (ThermoFisher Cat: 4392422) using Invitrogen Lipofectamine RNAiMAX Reagent (Thermofisher, Cat: 13778075). After 24 hours (UOK276) or 48 hours (RCJ), cells were lifted and re-seeded for various assays. For overexpression of Pfn1 in UOK276, a doxycycline inducible overexpression system was used. UOK276 overexpression cells and UOK276 wildtype cells (control) were treated with doxycycline (1 μg/ml) for 24 hours before replating for various assays. For pharmacological Profilin1 inhibition experiments, cells were plated for various assays with 0uM, 100uM, 200uM Profilin1 inhibitor C74 with DMSO as a vehicle control.

### Western Blot

Cells were lysed at end point experiments with IP buffer containing IP buffer (1 M Tris HCl pH 7.5 (25 mM), 5 M NaCl (150 mM), 0.5 M EDTA (1 mM), Triton X 100 (1%), glycerol (5%))) and 1% SDS. Cells were scraped and lysates were centrifuged. Protein was collected and stored at -80°C until further use. Protein was mixed with 6x Laemmli SDS sample buffer (6x, Reducing) and boiled for 5 min. Proteins were separated on 4-20% polyacrylamide gel (Invitrogen, XP04202BOX) in MOPS buffer (50.4 mM Tris, 50.2 mM MOPS, 3.47 mM SDS, 1.03 mM EDTA) and transferred onto a Trans-Blot Turbo Transfer Membrane (Bio-Rad). Samples were blocked with 10% milk in TBS and primary antibodies were diluted in 10% milk in TBS-T. Samples were incubated with antibodies Profilin1 (Novus, EPR6304) and GAPDH (Sigma, G9545) for 1 hour at room temperature on a plate rocker, washed in TBS-T, and incubated with secondary HRP conjugated antibody at room temperature for 1 hour. Signal was developed using Clarity Western ECL Substrate (BioRad, 1705060) and imaged. Protein abundance was quantified by densitometry and normalized to GAPDH. Values were then expressed relative to the control group.

### Scratch Assay

24 hours after transfection or induction of overexpression, UOK cells were plated in 96 well dish at 50,000 cells/well and grown overnight. Confluent wells were scratched with a 10uL pipette tip and imaged immediately and again 24 hours later. Distance of ingrowth of scratch was measured using ImageJ. Values were normalized to the averaged control well distances.

### Proliferation Assay

24 hours (UOK) or 48 hours (RCJ) after transfection or overexpression, cells were replated in a 24 well dish at 10,000 cells/well. Cells were left for 4 days with media replaced on day 2. For C74 proliferation assays, cells were seeded in media supplemented with 0uM, 100uM, 200uM Profilin1 inhibitor C74 with DMSO as a vehicle control. Media was replaced on day 3. Cells were lifted from the plates and counted, and cell counts were normalized to the averaged control well cell counts.

### Immunofluorescence Imaging

24 hours after transfection or overexpression induction, UOKs were plated on a glass coverslip coated with 50ug/mL type I collagen in a 24 well plate and left for 48 hours. Cells were fixed in 3.5% PFA for 5 minutes at 37°C. Coverslips were then washed twice with PBS and stored in 4°C fridge. For immunofluorescence, cells were permeabilized with 0.5% triton x for 5 minutes. Cells were blocked for 15 minutes in 10% goat serum in phosphate buffered saline (PBS). Cells were incubated with primary antibodies TOM20 (1:200) (Invitrogen Cat: 11802-1-AP) and tubulin (1:200) (Invitrogen Cat: PA5-21979) for 1 hour and washed with 0.02% PBS-tween. Cells were then incubated with secondary antibodies anti-rabbit FITC (1:200) (Jackson Immunoresearch Cat: 111-095-003) and anti-mouse far red (1:200) (Jackson Immunoresearch Cat: 115-605-116) and counterstained with rhodamine phalloidin and DAPI for 1 hour. Coverslips were rinsed twice with PBS-tween, once with PBS, and once with water. Coverslips were mounted on slides and left to dry overnight before imaging. Slides were imaged on the Olympus BX61. 5 images per slide were captured. TOM20 was quantified using CellProfiler. TOM20 staining was used to assess mitochondrial connectivity through tortuosity and branch length measurement. Individual cell mitochondrial analysis employed ImageJ skeletonization and quantifi:ion of branch lengths. Tortuosity was calculated as branch length/Euclidean distance.

### Colony Forming Assay

24 hours (UOK) or 48 hours (RCJ) after transfection or overexpression, cells were replated on BME (R&d Systems Cat: 3445-005-01) in a 96 well dish at 5×10^3^ cells/well. Wells were imaged 24 hours later (day 1), day 2 and/or 3, and day 4 and/or 5. Each day’s data was analyzed using an unpaired student t-test, normalized to control wells. Data was binned as day 1, day 2-3, and days 4-5. For C74 proliferation assays, cells were seeded in media supplemented with 0uM, 100uM, 200uM Profilin1 inhibitor C74 with DMSO as a vehicle control. Media was replaced on day 3.

### Transwell Invasion Assay

24 hours (UOK) or 48 hours (RCJ) after transfection or overexpression, cells were replated in a 24 well plate with transwell inserts. Cells were plated in serum free DMEM at 7.5×10^4^ cells/well for KD and at 5×10^4^ cells/well for overexpression cells. The bottom of the well was filled with 0.5 ml respective cell line media with 10% FBS. The inserts were fixed 48 hours later in 3.5% PFA for 5 minutes at 37°C. The inserts were then washed twice with PBS, and the tops of the inserts were scraped with cotton tip applicators and washed with PBS again. The inserts were then incubated in a 0.1% crystal violet staining solution for 20 minutes at room temperature and then washed 3 times with PBS and 1 time with DI H2O. The inserts were allowed to dry overnight before imaging. 5 images per well were taken and ImageJ was used to quantify migrated cells through quantification of % area invaded, normalized to control wells.

### Edu Assay

24 hours after transfection or overexpression, UOK276 cells were replated on a collagen I coated glass coverslip in a 24 well plate and incubated overnight. Using Click-iT® EDU Imaging Kit (ThermoFisher Cat: C10340) cells were incubated with 10uM EDU solution for 2 hours and then fixed with 3.7% PFA for 5 min at 37°C. Coverslips were washed with 3% BSA in PBS in PBS and stored in 20°C until further use. On the day of EDU detection, the assay kit directions were followed, and the cells were additionally counterstained with Rhodamine/Phalloidin 535 and Hoechst stain for 20 minutes in the dark. Coverslips were rinsed twice with PBS-tween, once with PBS, and once with water. Coverslips were mounted on slides and left to dry overnight before imaging. Slides were imaged on the Olympus BX61. Total and % EDU positive cells were calculated using CellProfiler and data was normalized to control wells.

### Seahorse Metabolic Flux Assay

24 hours after transfection, UOK276 cells were plated in a 8-well Agilent Seahorse XF Cell Culture Microplate in UOK276 standard media. Another 24 hours later, the media was changed with XF DMEM supplemented media, and manufacturer’s protocol was followed for the Agilent Seahorse XFp Cell Mito Stress Test Kit (Agilent Technologies, 03015-100).

### JC1 and CellROX

24 hours after transfection or overexpression induction, UOKs were replated on a glass 8 well plate coated with type I collagen and incubated for 48 hours. Cells were then incubated with JC1 (1:2000) (Invitrogen Cat: M34152A) and CellROX (1:500) (Invitrogen Cat: c10422) for 30 minutes at 37°C. Wells were imaged immediately at 5 images per well at 60x magnification on the Olympus BX61. JC1 and CellROX intensity were measured using ImageJ integrated density measurement tool. Data was normalized to control group intensities.

### Statistics

A Student’s t-test was used to assess statistical significance on the normalized values of control and knockdown biological replicates. In proliferation, migration, invasion, imaging assays, 2 technical replicates were used across 3 biological replicates. In colony forming and seahorse assays, 3 technical replicates were used across 3 biological replates. A one-way ANOVA was used to determine significance in the proliferation and colony assays with varying C74 treatments.

## RESULTS

### Pfn1 expression increases in stage IV ChRCC

Analysis of TCGA KICH dataset showed that at stage IV of ChRCC, patients have significantly increased Pfn1 expression (**Fig 1A**), though it is unknown if Pfn1 is a reason for or byproduct of the advanced disease. GEPIA analysis of TCGA data revealed when ChRCC patients were stratified between high and low Pfn1 expression, those with high Pfn1 expression had a shorter survival time (**Fig 1B**) (HR = 4.2, log-rank p=0.053), suggesting a potential association with advanced disease progression and ChRCC aggressiveness. Together these data implicate Pfn1 in advanced ChRCC and provided the rationale for functional investigation.

**Figure 1:**
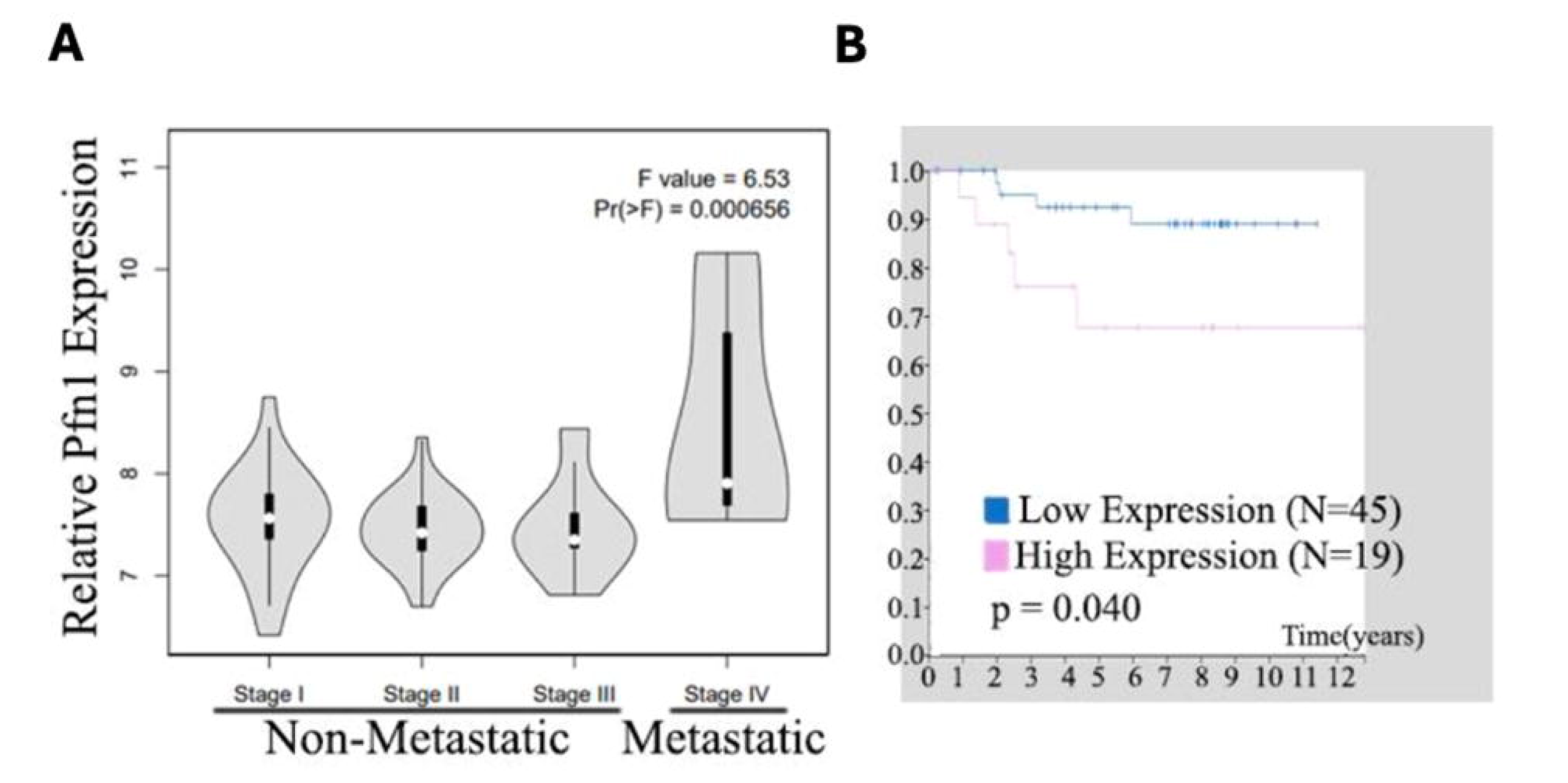
Pfn1 expression associates with advanced disease stage and poor survival in ChRCC. (**A**) Pfn1 expression at increasing stages of ChRCC from N=66 patient samples. Data was plotted on Gene Expression Profiling Interactive Analysis (GEPIA) platform (**B**) Kaplan-Meier survival data of patients with ChRCC and Pfn1 expression. source: Human Protein Atlas. P-value=0.040.

### Loss of Pfn1 decreases ChRCC cell proliferation and motility

To determine the functional consequences of Pfn1 loss in ChRCC, we performed siRNA-mediated Pfn1 knockdown in two ChRCC cell lines, UOK276 and RCJ41M. Western blot analysis confirmed significant reduction in Pfn1 protein levels in knockdown cells relative to scrambled siRNA controls in both UOK276 (**Fig. 2A, B**) and RCJ41M (**Fig. 2A, C**). Following knockdown, cell accumulation over four days was significantly reduced in both lines, with a 55% decrease in UOK276 (**Fig. 2D**) and a 29% decrease in RCJ41M (**Fig. 2E**) relative to controls. To determine whether this reduction reflected decreased proliferation, we conducted an EdU assay on UOK276 cells and found no difference between control and Pfn1 KD cells (**Fig. 2F**). However, trypan blue exclusion at the four-day endpoint demonstrated a significant 20% reduction in the proportion of live UOK276 cells in the Pfn1 knockdown condition (**Fig. 2G**), suggesting that increased cell death, underlies the decrease in cell accumulation. We also evaluated the impact of loss of Pfn1 in RCJ41T2 cell line, derived from the primary tumor of the same patient as RCJ41M cell line (which is from a metastatic lesion from that patient) was also tested. Interestingly, RCJ 41T2 Pfn1 KD cells did not have a significant decrease in cell number compared to control, although trends indicate a slightly lower cell count compared to control (**Supp. Fig 1A-B**), suggesting that there may be a more significant role in the metastatic variant of the RCJ line.

**Figure 2:**
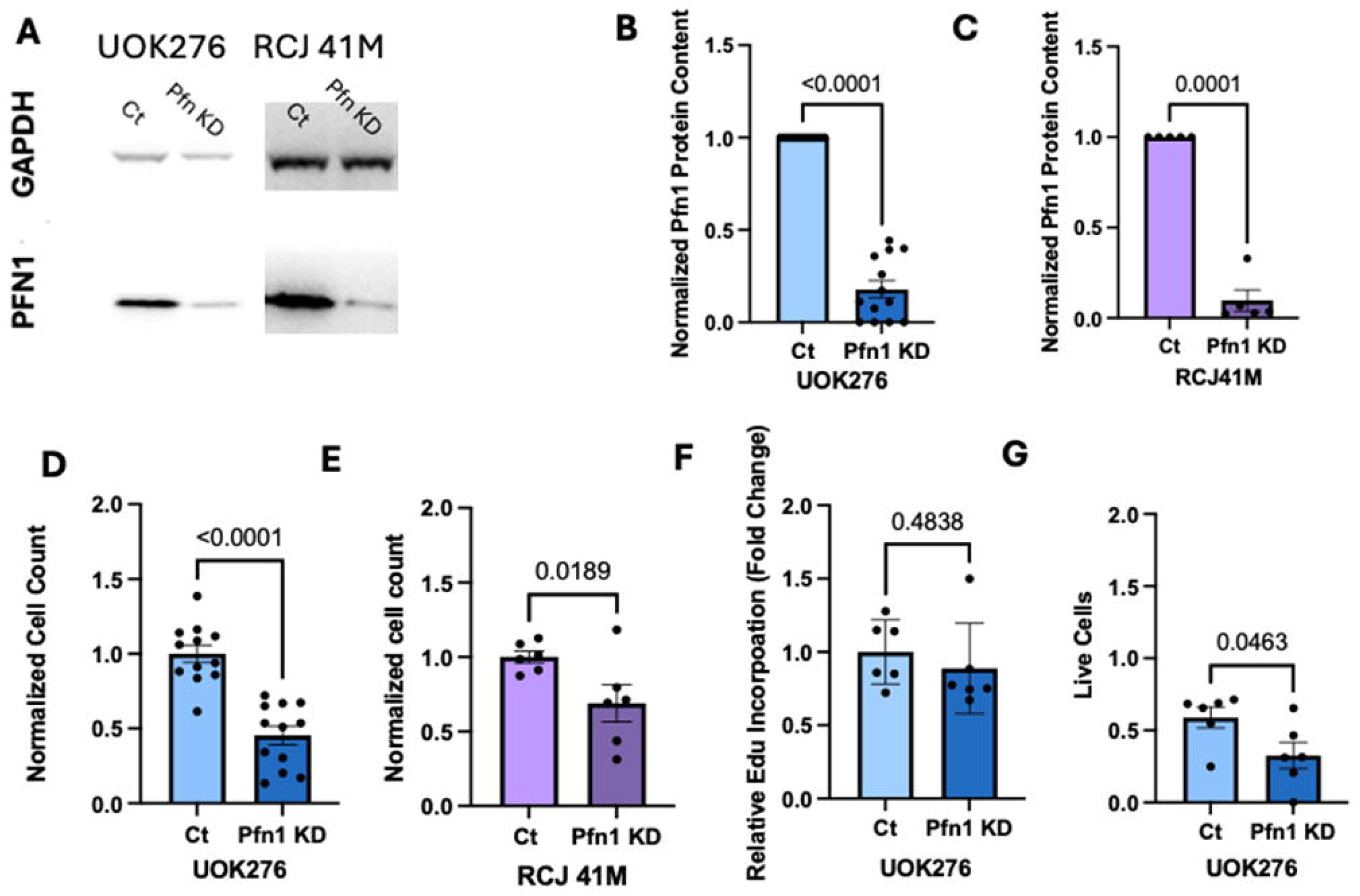
Pfn1 knockdown reduces cell accumulation and viability in ChRCC cell lines. Western blot validation of (**A**) Pfn1 knockdown (KD) in UOK276 (n=13) and RCJ41M (n=5) Student’s unpaired t-test. Pfn1 protein content was normalized to GAPDH through densitometry in (**B**) UOK276 and (**C**) RCJ41M. Values were then normalized relative to control groups, Student’s unpaired t-test. Normalized cell number after 4 days of culture in Pfn1 knockdown in (**D**) UOK276 (n=5) and (**E**) RCJ41M (n=3), Student’s unpaired t-test. (**F**) Normalized Edu incorporation quantified from images in UOK276 cells, n=3, Student’s unpaired t-test. (**G**) % live UOK276 cells after 4 days proliferation, student’s unpaired t-test (n=3).

### Downregulation of Pfn1 depletes chromophobe tumorigenic abilities

To understand if Pfn1 loss reduces the metastatic/severity level of ChRCC, colony forming and invasion assays were conducted. Colony forming assays were conducted. UOK276 and RCJ41M had significant impairments of colony formation with Pfn1 knockdown. UOK276 had a 70% reduction in colony area at day 4 compared to control cells (**Fig. 3A, B**) and RCJ41M 50% reduction at day 4 (**Fig. 3C, D**). To assess functional migratory/invasion ability with Pfn1 depletion, transwell invasion assays were conducted. Transwell invasion assays similarly revealed reduced invasive capacity in Pfn1 knockdown cells, with UOK276 showing a 33% decrease (**Fig. 3E, F**) and RCJ41M a 75% decrease in invaded area relative to controls (**Fig. 3G, H**). We also performed a scratch assay with UOK276 as an orthogonal approach and found that Pfn1 KD cells had a 45% reduction in gap closure compared to control cells (**Supp. Fig 2**). Consistent with previous findings, RCJ41T2 cells were not as greatly affected by siRNA mediated pfn1 knockdown, with no difference found between controls and pfn1 knockdown in % area of colonies (**Supp. Fig 3A)** nor the % area invaded in the transwell assay (**Supp. Fig3B**).

**Figure 3:**
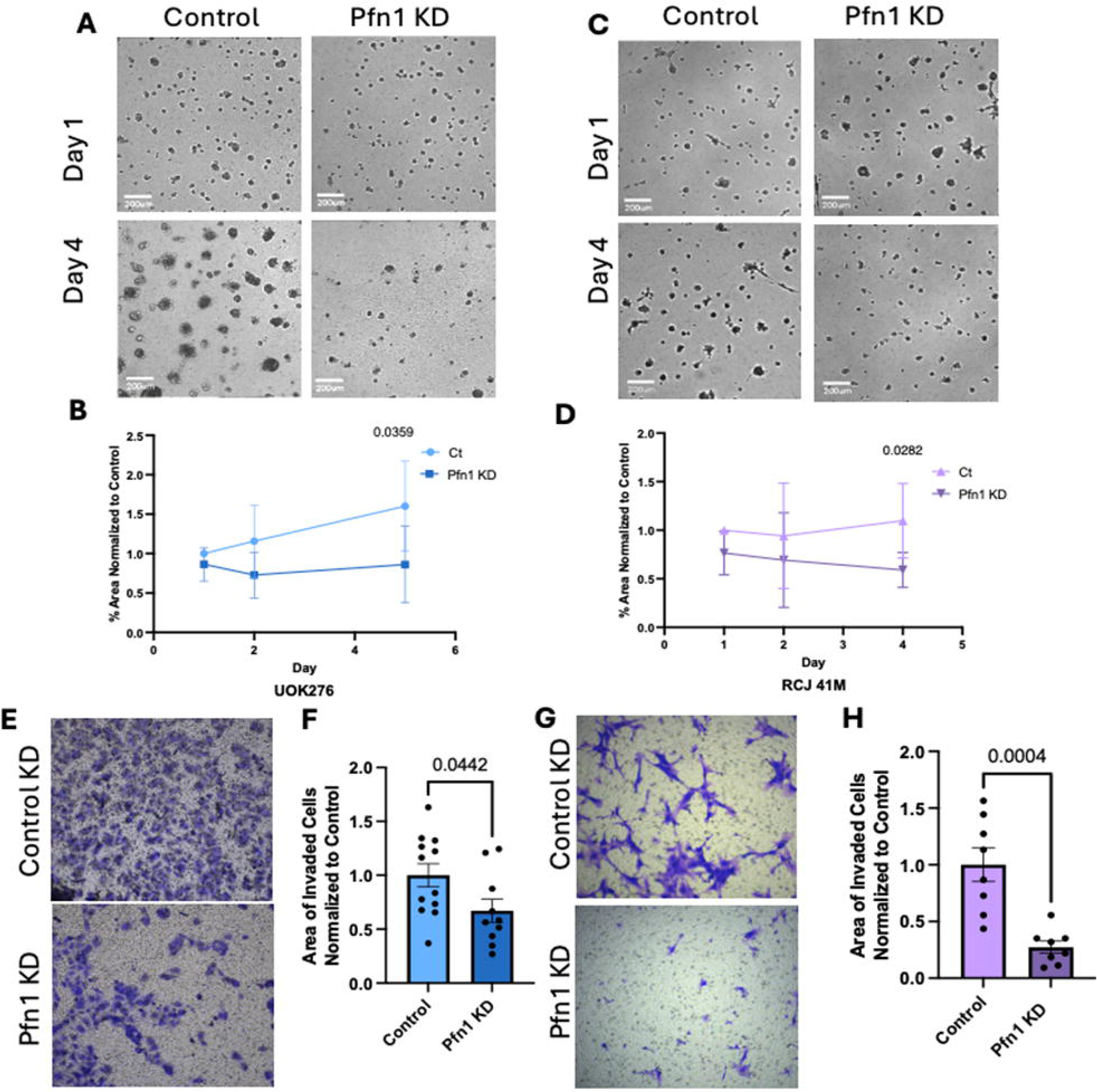
Pfn1 knockdown impairs colony formation and invasion in ChRCC cell lines. Representative colony forming assay images at Day 1 and Day 4 in (**A**,**B**) UOK276 Pfn1 KD cells (n=5) and (**C**,**D**) RCJ41M Pfn1 KD cells (n=3), student t-test. Scale bars = 200µm. Quantification of % colony area normalized to control area at Day 1 (t=24 hours), Day 2-3 (t=48-72 hours), and Day 4-5 (96-120 hours). Representative images of crystal violet stained (**E, F**) UOK276 (n=6) and (**G, H**) RCJ41M (n=3) after a 24 and 48-hour transwell invasion assay respectively in Pfn1 KD cells. Cells were fixed and stained with crystal violet, student t-test.

### Overexpression of Pfn1 enhances UOK276 proliferation, colony forming, and invasion abilities

To test whether the phenotypic and functional effects in Pfn1 knockdown cells could be not only inversed but enhanced, Pfn1 overexpression was conducted in UOK276. Pfn1 overexpression (OE) using a doxycycline inducible overexpression was verified by western blot in control and Pfn1 OE UOK276 cells (**Fig. 4A, B**). Pfn1 overexpression resulted in a 60% increase in cell accumulation over four days compared to controls (**Fig. 4C**), mirroring the reciprocal effect observed with knockdown. Scratch wound assays revealed an 180% greater gap closure in overexpression cells relative to controls at 24 hours (**Fig. 4D, E**). EdU incorporation was unchanged between overexpression and control cells (**Fig. 4F**), consistent with the knockdown findings and suggesting that Pfn1 modulates cell survival and motility rather than proliferative rate per se in this context. Transwell invasion assays demonstrated a fourfold increase in invaded area in Pfn1 overexpression cells compared to controls (**Fig. 4G, H**). Colony forming assays showed no statistically significant difference in colony area between overexpression and control cells at days 4–5, though a trend toward accelerated early colony growth was observed at days 2–3 (**Fig. 4I**). Together, these data establish a bidirectional relationship between Pfn1 expression and ChRCC aggressive phenotypes, with Pfn1 loss attenuating and Pfn1 gain enhancing migratory, invasive, and survival-associated behaviors.

**Figure 4:**
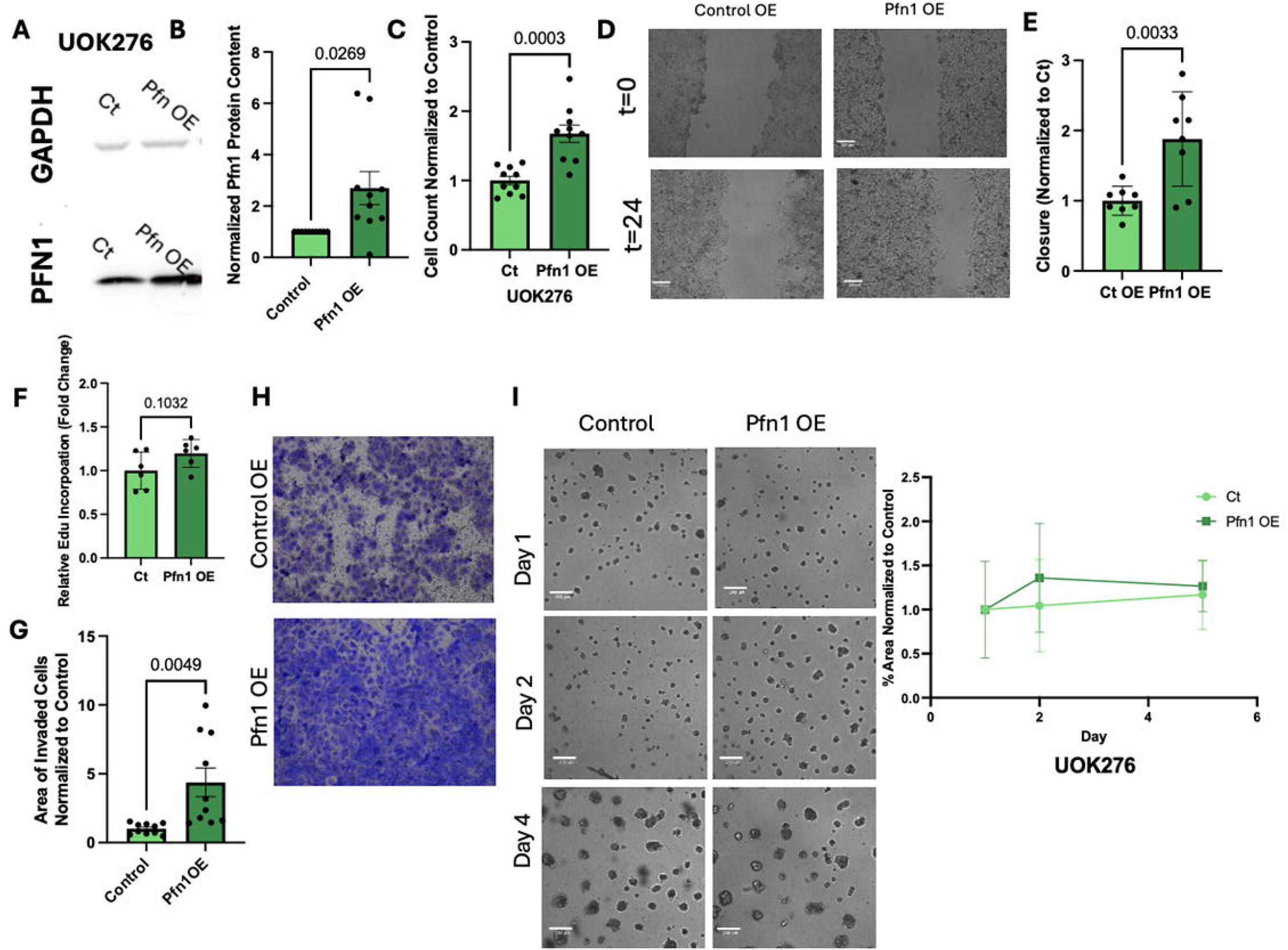
Pfn1 overexpression enhances cell accumulation, migration, and invasion in UOK276 cells. Western blot validation of Pfn1 overexpression (OE) (n=10) in UOK276 cells, student t-test (**A**,**B**). (**C**) Normalized cell number after 4 days of culture in Pfn1 overexpression cells normalized to control cells, n=5, Student’s unpaired t-test. (**D**) Representative scratch and closure images at timepoint (t) = 0 and 24 hours in UOK276 Pfn1 overexpression cells. Scale bars = 200µm. (**E**) Quantification of % closure in Pfn1 overexpression normalized to control. Distance of scratches were measured at time of scratch (t=0) and 24 hours later (t=24). n=3, Student’s unpaired t-test. (**F**) Normalized Edu incorporation quantified from images in UOK276 Pfn1 OE cells, n=3, Student’s unpaired t-test. (**G**) Representative images of crystal violet stained UOK276 Pfn1 overexpression cells (n=4) after a 48-hour transwell invasion assay. Cells were fixed and stained at t=48 hours, Student’s unpaired t-test. Scale bars = 200µm. (**H**) Quantified % area of crystal violet stained cells, Student’s unpaired t-test. (**I**) Colony forming assay images and quantification at Day 1 (t=24 hours), Day 2-3 (t=48-72 hours), and Day 4-5 (96-120 hours) in UOK276 Pfn1 OE cells (n=5), Student’s unpaired t-test. Scale bars = 200µm.

### ROS accumulation inversely correlates with Pfn1 knockdown and overexpression in UOK276 cells

Given ChRCC’s established mitochondrial dysfunction and sensitivity to oxidative stress, we investigated whether Pfn1 modulation influences intracellular reactive oxygen species (ROS) levels in UOK276 cells. CellROX fluorescence intensity was significantly increased by 2.3-fold in Pfn1 knockdown cells relative to scrambled controls, while overexpression reduced CellROX signal by 57% (**Fig. 5A-B**). These findings demonstrate that Pfn1 expression regulates intracellular ROS in ChRCC cells, with loss of Pfn1 driving oxidative stress accumulation and gain of Pfn1 suppressing it. This pattern is consistent with the cell accumulation and viability data, raising the possibility that ROS-mediated cytotoxicity contributes to the reduced cell survival observed with Pfn1 knockdown. Assessment of mitochondrial membrane potential using the JC-1 probe revealed no significant difference between Pfn1 knockdown and control cells; however, Pfn1 overexpression was associated with a significant 40% reduction in JC-1 red channel fluorescence intensity (**Fig. 5C-D**), suggesting that overexpression of Pfn1 modestly perturbs mitochondrial membrane potential in this context. Mitochondrial morphology, assessed by TOM20 immunofluorescence, was unaffected by either Pfn1 knockdown or overexpression (**Supp. Fig 4A-D**). To elucidate how Pfn1 may be affecting ROS, we explored several pathways. The Agilent Seahorse Mito Stress Test metabolic flux assay was used to assess mitochondrial function in Pfn1 KD cells and OE cells. This test uses oligomycin, FCCP, and Rot/A to assess mitochondria’s ability to withstand perturbation and recover. The data overall shows unreactive mitochondria, but an overall trend of reduced oxygen consumption rates (OCR) in Pfn1 KD compared to control, though statistically insignificant. Though the overexpression cells had an on average higher OCR compared to control groups, the Pfn1 OE cells did not respond to the various drugs in the mito stress test kit, suggesting these cells also harbor unhealthy mitochondria (**Supp. Fig 4E**). The Seahorse assays suggest Pfn1 loss or gain affects mitochondrial function in ChRCC cells but our results did not reach statistical significance. Ferroptosis, an emerging vulnerability in ChRCC^4,21–23^, was investigated by measuring glutathione levels. No differences were seen in glutathione accumulation in Pfn1 knockdown or overexpression UOK cells (**Supp. Fig 4F**).

**Figure 5:**
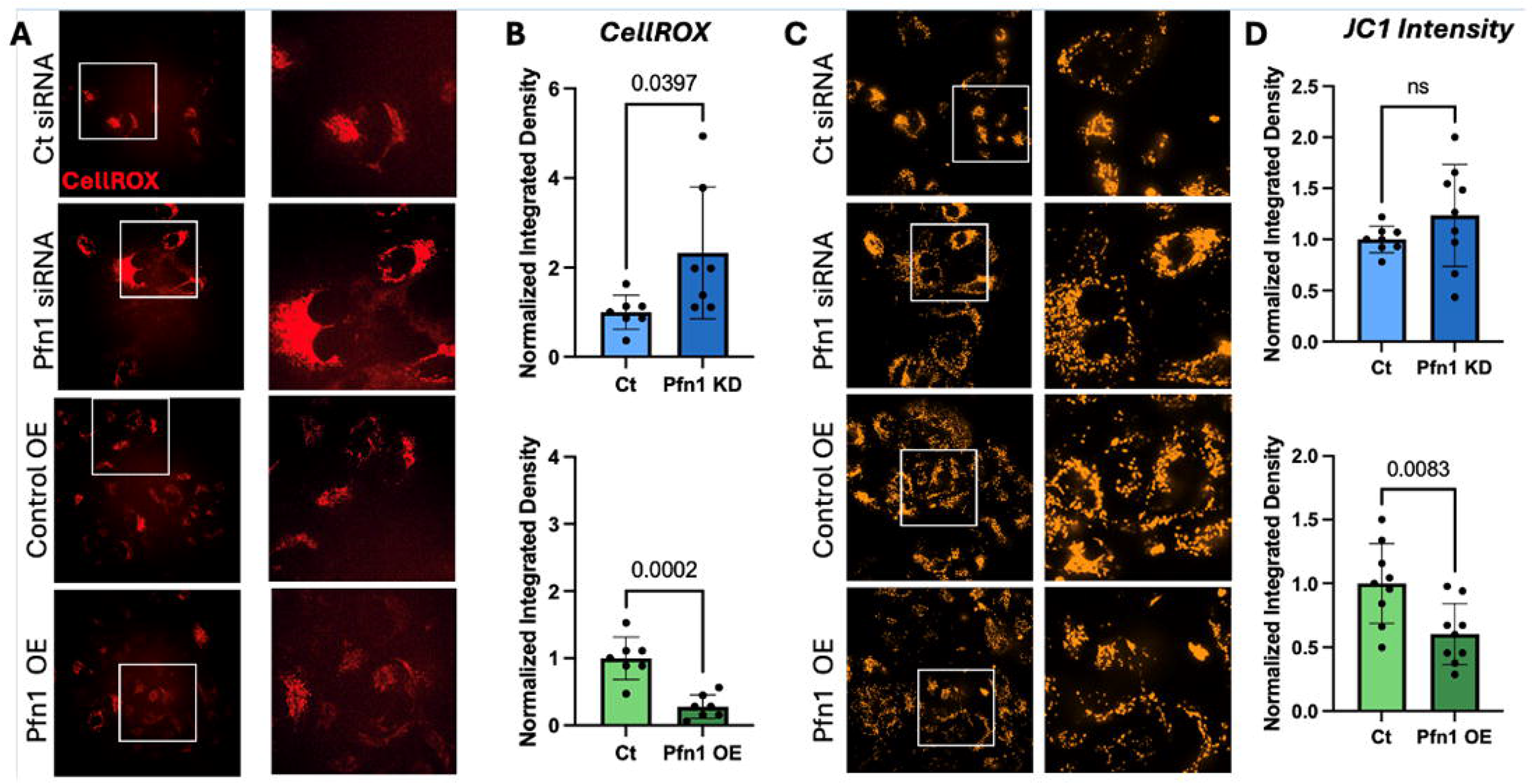
Pfn1 modulation bidirectionally regulates intracellular ROS in UOK276 cells. (**A**) 60x images and (**B**) quantification of CellROX intensity in UOK276 Pfn1KD and Pfn1 OE cells normalized to control intensity, Student’s unpaired t-test, n=3. (**C**) 60x images of UOK276 cells treated with JC1 in control, Pfn1 KD, and Pfn1 OE cells. (**D**) Quantification of JC1 intensity in UOK276 Pfn1KD and Pfn1 OE cells normalized to control intensity, Student’s unpaired t-test, n=3.

### Pharmacological inhibition of Pfn1 suppresses proliferation and colony formation across ChRCC cell lines

To extend genetic findings toward therapeutic relevance, we used C74^15,24^ (a small molecule inhibitor that blocks the Pfn1-actin interaction, across all three ChRCC cell lines at 0, 100, and 200µM. Treatment with 200µM C74 resulted in a significant reduction in cell number after 4 days of culture in all 3 cell lines. Cell counts were reduced by 45% in UOK276 (**Fig. 6A**) and 85% in RCJ41M (**Fig. 6D**). Colony forming ability was assessed over 5 days and % area colony was quantified. At day 4-5, UOK276 cells treated with 200µM C74 had a 1.5-fold decrease in colony area (**Fig 6B, C**) and RCJ41M cells treated with 200µM C74 had an approximately 90% reduction in colony area (**Fig 6 E, F**). Cell counts were reduced with 100 and 200µM C74 by 45% in and 90% respectively in RCJ41T2 (**Supp. Fig 5A**). While colony area was not significantly reduced at either dose, treated cells exhibited marked morphological disruption including loss of the branching, cord-like colony structures characteristic of control wells (**Supp. Fig 5B, C**). Since pharmacological Pfn1 inhibition affected the cells in a similar way as siRNA-mediated knockdown, this suggests that disruption of Pfn1 could be a putative therapy for ChRCC.

**Figure 6:**
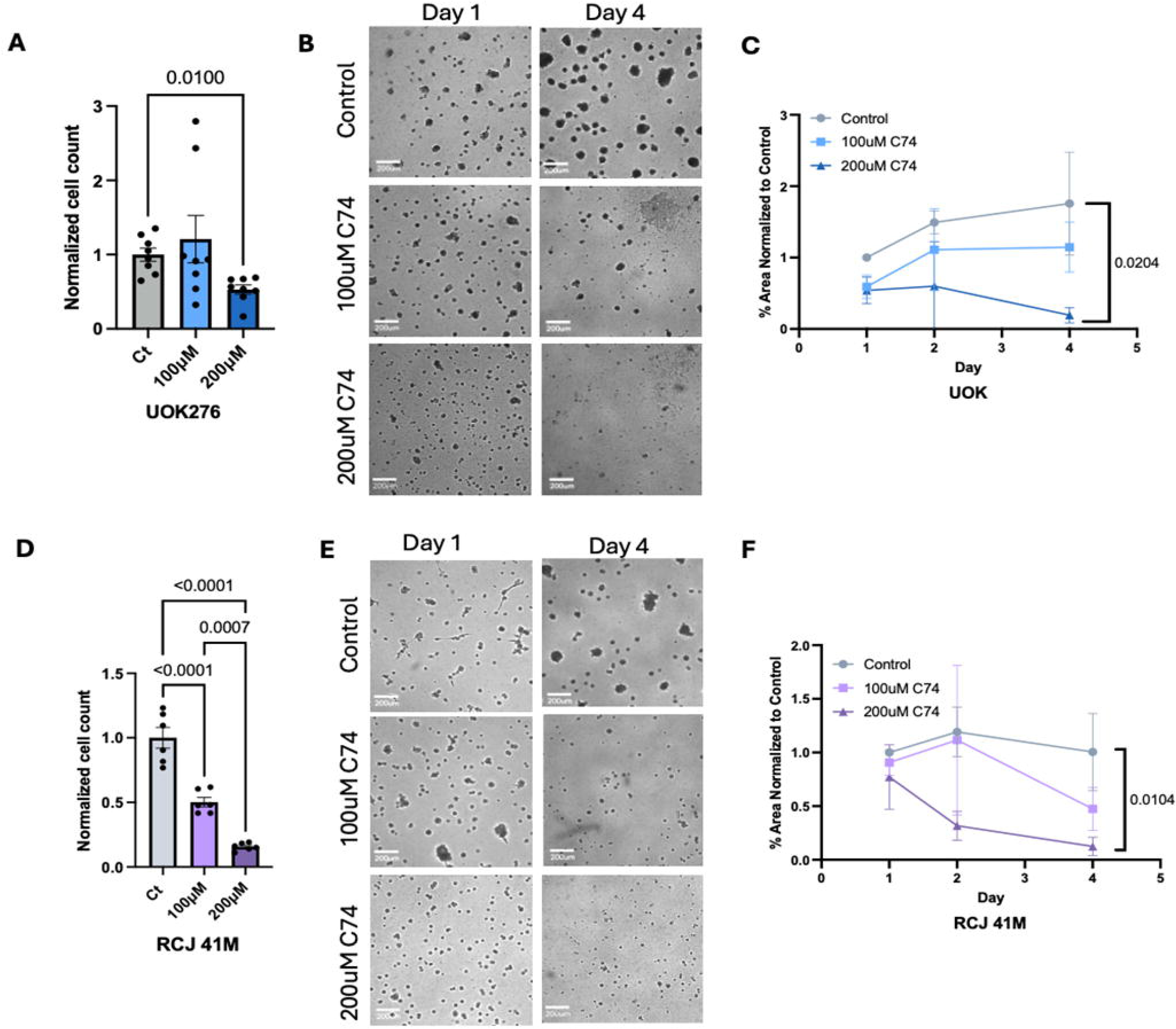
Pharmacological inhibition of Pfn1 with C74 suppresses proliferation and colony formation in ChRCC cell lines. Proliferation and colony forming assays in chromophobe cell lines with 0, 100, and 200µM C74 treatment. 4-day proliferation assay in (**A**) UOK276 (n=4) and (**D**) RCJ 41M (n=3), one-way ANOVA. Representative images of (**B**) UOK276 (n=4) and (**E**) RCJ 41M (n=3) and quantified % area of colony forming assay in (**C**) UOK276 (n=4) and (**F**) RCJ 41M (n=3), one-way ANOVA at each timepoint (Day 1, Day 2, Day 4).

## DISCUSSION

The present study demonstrates that Profilin-1 (Pfn1) is a critical regulator of aggressive cellular behavior in chromophobe renal cell carcinoma. Using complementary genetic and pharmacological approaches across three ChRCC cell lines, we show that Pfn1 promotes cell survival, migration, invasion, and colony formation, and that its loss drives intracellular ROS accumulation. These findings show that Pfn1 is a contributor to ChRCC progression and identify Pfn1-actin interaction as a potential therapeutic vulnerability in this treatment-neglected disease.

To interrogate Pfn1’s function, we utilized both genetic and pharmacological strategies. Pfn1 knockdown in UOK276 and RCJ41M cells reduced migration, invasion, and colony formation. This data is consistent with Pfn1’s known role in actin dynamics and motility, as an impaired actin system would also impart decreased migratory abilities^12,25^. Pfn1 knockdown significantly reduced cell number, whereas overexpression significantly enhanced it. An EdU assay was conducted to determine whether this difference in cell number was due to actual proliferation in UOK276 cells. Interestingly, we found no difference in the EdU incorporation in both Pfn1 knockdown and overexpression cells. More compellingly, trypan blue exclusion at the four-day endpoint revealed a significant 20% reduction in live cell proportion in Pfn1 knockdown conditions, suggesting that increased cell death rather than reduced proliferative rate underlies the decrease in cell accumulation. This interpretation is further supported by the CellROX data where we show Pfn1 knockdown drove a 2.3-fold increase in intracellular ROS, while overexpression reduced ROS by 57%, establishing a relationship between Pfn1 expression and oxidative stress that mirrors the cell accumulation phenotype. Elevated ROS is a well-established inducer of cell death, and its accumulation following Pfn1 loss in ChRCC cells at least partially explains the reduced cell count. Several studies have demonstrated ChRCC’s fragile sensitivity to ferroptosis^21,22^, thus we explored if the reduced tumorigenic properties seen with Pfn1 KD is due to ferroptotic upregulation. We measured reduced glutathione levels, and again, found no difference between groups. While these data do not support ferroptosis as the primary mechanism downstream of Pfn1 loss in ChRCC, they do not preclude a role for other ROS-mediated death pathways, and further mechanistic dissection is warranted.

RCJ41T2 cells did not exhibit the same sensitivity to siRNA mediated Pfn1 knockdown. Importantly, RCJ41T2 and RCJ41M cell lines were derived from the same patient, with 41T2 originating from the primary tumor and 41M the metastatic lesion. This distinction may account for the differential sensitivity of the two cell lines. Our data suggesting the more advanced ChRCC tumors, the higher Pfn1 expression, may explain why RCJ41M are more susceptible to Pfn1 depletion compared to 41T2, again accentuating the therapeutic potential of Pfn1 inhibition in advanced ChRCC.

Previous studies have reported links between Pfn1 and mitochondrial regulation. Read et al. recently found Pfn1 loss led to enlarged and elongated mitochondria, increased mitophagy, and accumulation of Pfn1 within mitochondria^26^. Similarly, Sun et al. demonstrated that Pfn1 knockdown increased mitophagy regulators Pink1 and Parkin, promoting the polarization of macrophages toward an inflammatory M1 macrophage phenotype in lung adenocarcinoma^17^. Although mitophagy markers Pink1 and Parkin were not analyzed in the present study, our findings do not fully align with the previous reports. Specifically, Pfn1 knockdown in our model led to ROS accumulation whereas enhanced mitophagy would typically be expected to remove damaged mitochondria and remove oxidative stress.^27^ These discrepancies most likely reflect cell-type-specific differences in Pfn1-mitochondrial interactions, emphasizing that Pfn1 biology cannot be assumed to operate uniformly across tumor contexts.

The present study has several important limitations. All functional data were generated in vitro, and the in vivo relevance of Pfn1-driven aggressiveness in ChRCC remains to be established. The mechanistic basis of ROS accumulation following Pfn1 loss has not been formally dissected, and the relative contributions of increased cell death versus reduced proliferation to the knockdown phenotype require further clarification. Nevertheless, the convergent evidence from genetic loss-of-function, gain-of-function, and pharmacological inhibition experiments presented here establishes Pfn1 as a functional driver of ChRCC aggressiveness and identifies C74-mediated disruption of the Pfn1-actin interaction as a potential therapeutic strategy deserving of further preclinical development. Future work will prioritize in vivo validation and mechanistic characterization of the Pfn1-ROS axis in ChRCC, with the goal of advancing cytoskeletal targeting as a subtype-specific therapeutic approach for patients with advanced disease.

## Acknowledgements

The authors acknowledge funding from NIH grant R00CA267180 (Gau), DoD KC240157 (Gau), and NIH T32HL076124 (Montanari).

## Figure Legends

**Supplemental Fig 1: RCJ41T2 cell line is not significantly impacted by loss of Pfn1.** (**A**) Pfn1 knockdown (KD) in RCJ 41T2 (n=3) cells, Student’s unpaired t-test. Pfn1 protein content was normalized to GAPDH through densitometry. Values were then normalized relative to control groups, Student’s unpaired t-test. (**B**) Normalized cell number after 4 days of culture in Pfn1 knockdown in RCJ41T2 (n=3), Student’s unpaired t-test.

**Supplemental Fig 2: Scratch Assay in UOK276 cells.** Representative images and quantification of % closure in UOK276 Pfn1 knockdown cells. Distance of scratches were measured and images were taken at time of scratch (t=0) and 24 hours later (t=24). n=3, Student’s unpaired t-test. Scale bars = 200µm.

**Supplemental Fig 3: RCJ41T2 colony formation and invasion not affected by genetic Pfn1 loss.** (**A**) Representative colony forming assay images at Day 1 (t=24 hours), Day 2-3 (t=48-72 hours), and Day 4-5 (96-120 hours) in RCJ41T2 Pfn1 KD cells (n=3), student t-test. Scale bars = 200µm and quantification of % colony area normalized to control area at Day 1 (t=24 hours), Day 2-3 (t=48-72 hours), and Day 4-5 (96-120 hours). (**B**) RCJ41T2 (n=3) after a 48-hour transwell invasion assay respectively in Pfn1 KD cells. Cells were fixed and stained with crystal violet, student t-test.

**Supplemental Fig 4:** (**A**) 60x Tom20 staining representative images in UOK276 control scrambled siRNA and Pfn1 KD and (**B**) control OE and Pfn1 OE. Bars= 50μm (**C**,**D**) Quantification of Tom20 staining, graphed as mitochondria area/cell area, n=3, Student’s unpaired t-test. (**E**) Seahorse metabolic flux assays displaying UOK276 n=3 replicates of oxygen consumption rates in control vs pfn1 KD and control vs PFN1 OE cells. (**F**) Glutathione measurement using reduced glutathione measurement kit in UOK276 Pfn1 KD and Pfn1 OE cells.

**Supplemental Fig 5: RCJ41T2 cell count and colony formation enhanced sensitivity to pharmacologic Pfn1 inhibition.** (**A**) Proliferation assay with 0, 100, and 200µM C74 treatment. 4-day proliferation assay in RCJ 41T2 (n=3) cells, one-way ANOVA. (**B**) Representative images and (**C**) quantified % area of colony forming assay in RCJ 41T2 (n=3), one-way ANOVA at each timepoint (Day 1, Day 2-3, Day 4-5).

