## Supplementary figures and images for "Profilin-1 Promotes Chromophobe Renal Cell Carcinoma Malignancy"

### Supplemental Figure 1

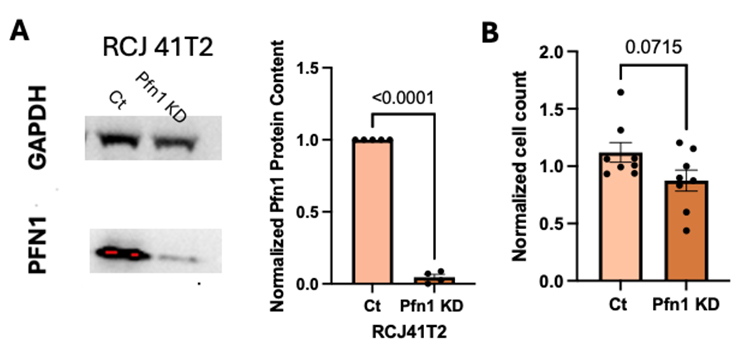

### Supplemental Figure 2

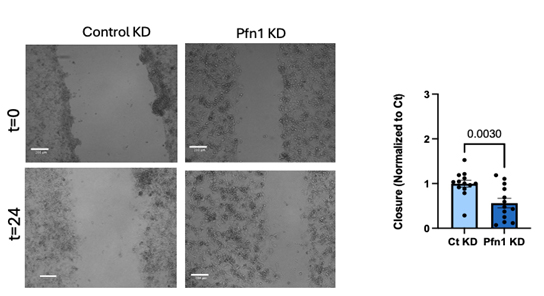

### Supplemental Figure 3

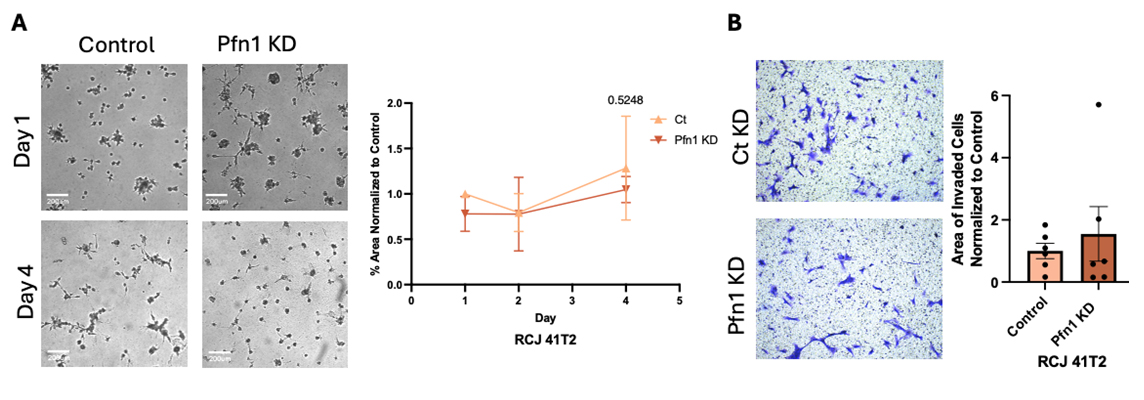

### Supplemental Figure 4

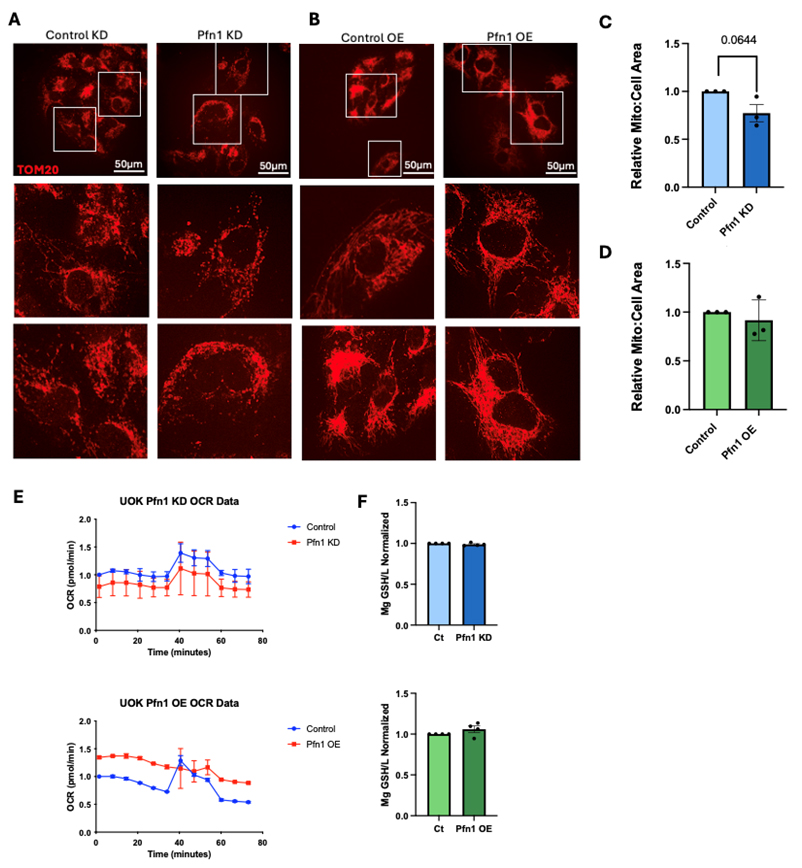

### Supplemental Figure 5

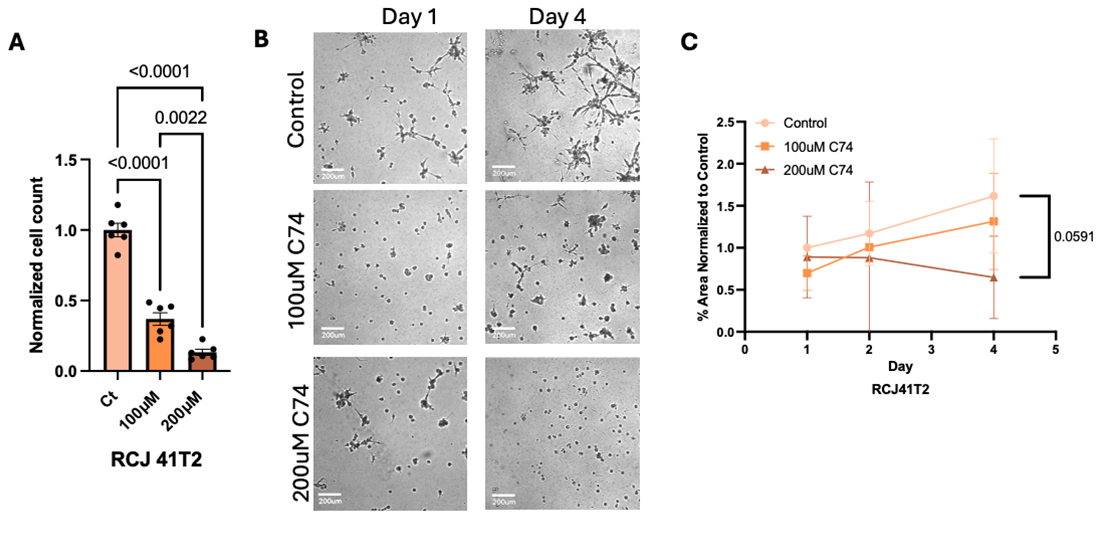
